# *Staphylococcus aureus* exhibits spatiotemporal heterogeneity in Sae activity during kidney abscess development

**DOI:** 10.1101/2025.07.08.663646

**Authors:** Anjali Anil, Rezia Era D. Braza, Bessie Liu, Valerie Altouma, Clement Adedeji, Ananyaa Welling, Alysha L. Ellison, Megan H. Check, Irnov Irnov, Rachel Prescott, Victor J. Torres, Kimberly M. Davis

## Abstract

Virulence factors are required for bacterial pathogens to establish infection, however, their expression can be energetically costly, and must be tightly controlled to avoid fitness costs. Expression can be controlled at specific stages during infection (temporal regulation) or expressed by small subsets of the bacterial population (spatial regulation). There has been a great deal of interest in developing virulence factor-targeting strategies to combat *Staphylococcus aureus* infection, but the spatiotemporal regulation of the virulence factor master regulatory systems (Agr, Sae) has not been explored during kidney abscess formation. This information is critical for the design of therapeutics targeting these pathways. Here, we utilized a fluorescent transcriptional reporter approach to visualize dynamics in Agr and Sae activity during abscess formation in the mouse kidney. We categorized kidney abscess formation into four stages, then defined spatiotemporal gene expression. Agr signalling appeared inactive in the kidney; consistent with this, *agr* mutant abscesses fully developed. In contrast, we observed heterogeneous (ON/OFF) activity of Sae at early stages where bacteria were found intracellularly within neutrophils. Sae activity increased as abscesses developed, and heterogeneity in spatial patterning was observed, but patterns varied between abscesses suggesting distinct microenvironments within individual abscesses.

Consistent with a requirement for Sae activity during abscess development, the *sae* mutant did not develop abscesses past early stages. These results have implications for the genes regulated by Agr and Sae, and suggest a requirement for Sae activity during kidney abscess development.

**Importance:** Infections with *Staphylococcus aureus* pose a serious public health threat due to high levels of antibiotic resistance and limited efficacy of alternative therapeutics. There has been a great deal of interest in developing novel therapeutics that target virulence factors essential during infection. However, it remains largely unknown if these factors are required at specific stages of the infection, and whether all bacterial cells or a limited subset express them. Here we sought to examine virulence factor expression using fluorescent reporter strains that would indicate activity of two master regulators of virulence in *S. aureus*, Agr and Sae. While Agr appeared inactive during kidney abscess development, the Sae system exhibited heterogeneity, increased expression at later stages, and was required for abscess progression. These results provide critical information for the development of virulence factor-targeting strategies for kidney abscess treatment.

## Introduction

Methicillin-resistant *Staphylococcus aureus* (MRSA) infections are a major cause of morbidity and mortality worldwide (1, 2), and therapeutic options can be scarce and limited in efficacy (3). Failure of vaccine or drug candidates in clinical trials has been largely attributed to the pathogen’s diverse array of virulence factors that subvert the host immune response (4). Neutralizing the effects of key virulence factors could curtail infection progression, making them attractive therapeutic targets (5). However, virulence factor expression can be energetically costly, and bacteria have adapted to express these gene products only at specific sites and stages of infection or within certain subsets of bacteria to minimize their fitness costs (6).

A hallmark of *S. aureus* systemic infection in mice is the formation of abscesses in the kidney. Abscesses consist of a central bacterial core, referred to as a staphylococcal abscess community (SAC), surrounded by necrotic and viable immune cells (7). Formation of an abscess sequesters bacteria, limiting further spread, yet clearance of the SAC by recruited immune cells is often unsuccessful and treatment requires antibiotic therapy, surgical intervention, or both (8).

Previous studies have shown a diverse array of virulence factors are necessary for abscess formation (9). Many of these virulence factors are regulated by the accessory gene regulatory system (Agr) and the *S. aureus* exoprotein expression (Sae) two component systems (10, 11). Agr is a quorum sensing system that enables *S. aureus* to modify gene expression as a function of its population density (12, 13).

Virulence factors expressed downstream of Agr include phenol-soluble modulins (PSMs), alpha-toxin, and certain toxins that are cytotoxic to immune cells (leukocidins). Sae can be activated in response to immune cell components and nutrient limitation (14–16), and promotes expression of leukocidins and other virulence factors previously shown to contribute to kidney abscess formation (9, 17). While virulence factors positively regulated by Agr and Sae are known to promote kidney abscess development, exactly when and in what bacterial subpopulations expression needs to occur is not well understood. The spatiotemporal expression patterns of *S. aureus* virulence factors are, therefore, an understudied component of abscess pathogenesis that could help inform a new generation of therapeutic approaches.

In this study, we utilized fluorescent reporter strains in the CA-MRSA USA300 LAC background in a mouse model of kidney infection to define spatiotemporal changes in the activities of Agr and Sae during kidney abscess development. The *agr* P2 promoter and the inducible *sae* P1 promoter were fused to *gfp* in distinct fluorescent reporter strains (15, 18). We found the Agr system was largely inactive during kidney abscess development, while the Sae system was active throughout abscess formation, with mature SACs showing heightened Sae activity. We also observed significant heterogeneity in Sae activity between bacterial subpopulations at different stages of infection. Using *agr* and *sae* mutant strains, we demonstrate that while Agr is dispensable, loss of Sae significantly impairs SAC development. Ultimately, our study underscores the significance of considering niche and stage specificity of virulence factor expression, and the presence of non-expressing bacterial subsets, in the development of virulence factor-targeting therapeutics.

## Results

### Retro-orbital inoculation results in gradual expansion of the *S. aureus* population in the mouse kidney

We hypothesized that *S. aureus* would exhibit spatiotemporal regulation of virulence factor expression during kidney abscess formation, and to assess virulence factor expression, we generated a series of fluorescent reporter strains. All strains contain a *P_sarA_*::*sod* RBS::*mCherry* cassette integrated at the SapI1 attachment site for constitutive expression of mCherry (19, 20). This allows mCherry signal to function as an internal control for the overall transcriptional activity of individual cells and allows us to quantify relative reporter expression normalized to mCherry signal. Transcriptional reporter constructs consist of the P2 promoter of the *agr* operon (*P_agrB_::gfp*) or the P1 promoter of the *sae* operon (*P_saeP_::gfp*) upstream of a *sod* ribosomal binding site (to ensure robust translation of the transcripts) fused to superfolder GFP (*gfp*). The GFP negative control strain (GFP^-^) has an insertion of a promoter-less version of *gfp*, which serves as both a baseline control for promoter-independent *gfp* expression, and detection of cellular and tissue-derived green autofluorescence. All *gfp* constructs were integrated into a neutral site in the genome of the mCherry^+^ strain (21).

We performed our initial experiments with the GFP^-^ control strain, employing a mouse model of systemic spread-derived kidney infection (22). We compared retro-orbital (RO) and tail vein (TV) inoculations, two commonly used routes for *S. aureus* systemic administration in mice (9, 23). TV inoculation with 10^6^ CFU of the GFP^-^ control strain resulted in a significantly higher bacterial load in the kidneys of mice at day 4 compared to day 2 post-inoculation (p.i.) (**Fig 1A**). The relatively high bacterial load at day 2 resulted in high levels of early tissue damage, early signs of morbidity, and suggested bacteria trafficked rapidly from the liver to the kidney instead of experiencing a bottleneck in the liver (23). For mice infected retro-orbitally (RO), the difference in kidney CFUs between days 2 and 4 p.i. was not statistically significant, indicating a slower increase in bacterial load (**Fig 1A**). For RO infection, the bacterial load continued to increase until day 5 p.i., suggesting initial seeding of kidneys and bacterial population expansion within the day 2 to 5 timeframe, beyond which a gradual decline suggests some control of infection (**Fig 1A**). Hence, for all subsequent experiments, we employed the RO route, which would allow us to monitor the entire range of kidney abscess dynamics (seeding, development, control). We also compared the bacterial load in the left and right kidneys of RO-inoculated mice at day 4 p.i. and observed no significant difference (**Fig 1B**). However, 2 of 7 mice had lower bacterial load and >10^4^-fold difference in CFUs between the two kidneys. For consistency, the left kidney was used for CFU enumeration and right kidney was used for microscopic analysis in all subsequent experiments.

**Figure 1:**
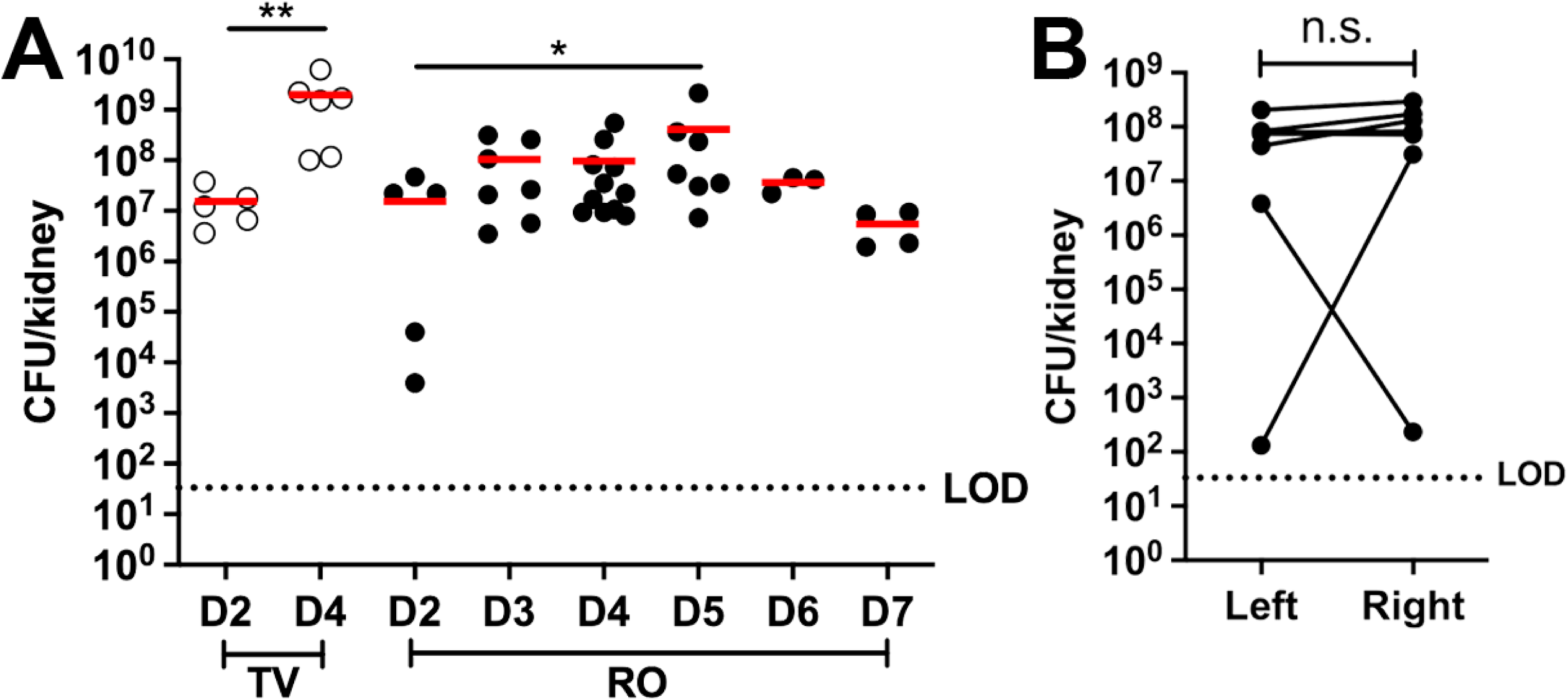
Retro-orbital inoculation results in gradual expansion of the *S. aureus* population in the mouse kidney. C57BL/6 mice were intravenously inoculated with 10^6^ CFU of the *S. aureus* GFP^-^ control strain (constitutive mCherry: *P_sarA_*::*sod* RBS::*mCherry*) via the tail vein (TV) or retro-orbital (RO) route. A) Bacterial load (CFU/kidney) at the indicated timepoints (D=day). Each dot represents one mouse. N = 3 to 11 mice per timepoint. Red bars: mean. B) Comparison of bacterial load in the right and left kidneys at D4 after RO inoculation. N = 7 mice. Lines connect pairs of kidneys from the same mouse. Horizontal dotted line: limit of detection (LOD). Statistics: A) Mann-Whitney t-test; B) Wilcoxon matched-pairs test. \*\**p*<0.01, \**p*<0.05, n.s.: not significant.

### Defining the four observed stages of *S. aureus* mouse kidney abscess development

We next sought to visualize and define the stages of *S. aureus* strain LAC kidney abscess development by fluorescence microscopy using the published framework with *S. aureus* strain Newman as a guide (9). C57BL/6 mice were inoculated RO with the GFP^-^ control strain, kidneys were harvested on days 3 through 5 p.i., and tissue was prepared for immunofluorescence microscopy. Analysis of kidney sections revealed distinct stages of abscess development which we broadly categorized into 4 stages, with both similarities and differences relative to previous studies (9). Stage 1 was defined by the presence of intracellular *S. aureus* within neutrophils (**Fig 2A, 2B**). This was consistent with the study by Pollitt et al. that demonstrated neutrophils play an important role in dissemination of *S. aureus* into the kidney (23), but this definition is distinct from the bloodstream stage 1 designation in Cheng et al. (24). Stage 2 was defined as small extracellular clusters of *S. aureus*, ranging in area between 4.03 to 16.07 µm^2^ (**Fig 2A, 2C**). A fully-formed, intact, SAC was categorized as stage 3, similar to previous designations (24). Most stage 3 SACs had a space (frequently autofluorescent) between bacterial cells and the surrounding host cells, pointing to the presence of the fibrin pseudocapsule or microcolony-associated meshwork (MAM) that serves as a physical barrier from surrounding host cells (17, 25). The area of stage 3 SACs ranged from 38.49 to 3129.52 µm^2^, with many SACs in the 50-1000 µm^2^ range (**Fig 2C**). We observed some SACs were disrupted, with host cells physically interacting with *S. aureus* at multiple points on the SAC; this was classified as stage 4 (**Fig 2A**).

**Figure 2:**
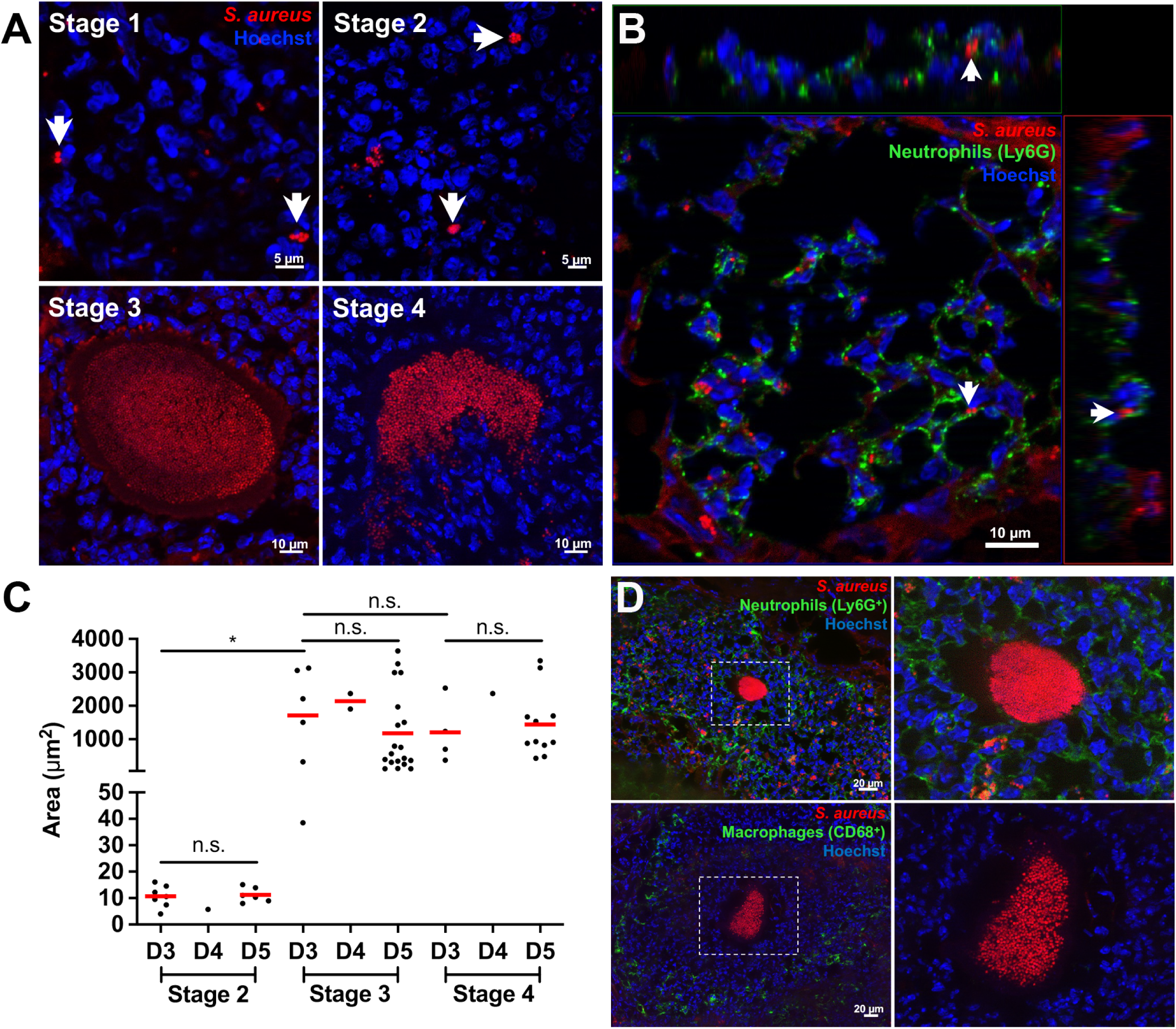
Defining the four observed stages of *S. aureus* mouse kidney abscess development. C57BL/6 mice were inoculated RO with the *S. aureus* GFP^-^ control strain. At the indicated timepoints (day 3 (D3) to day 5 (D5)), mice were sacrificed, and kidneys were harvested. A) Representative images depicting stages of S. *aureus* kidney abscess development. Stage 1: intracellular *S. aureus* (white arrow), stage 2: small extracellular clusters (white arrow), stage 3: fully formed and intact staphylococcal abscess community (SAC), and stage 4: dispersed SAC. B) Immunofluorescence images showing *S. aureus* inside neutrophils (Ly6G^+^). XZ and YZ planes corresponding to the point highlighted by the white arrow are shown at the top and right, respectively. C) Areas of *S. aureus* extracellular clusters (stage 2) or SACs (stages 3 and 4) at D3 to D5. Dots represent individual extracellular clusters (stage 2) or SACs (stages 3 and 4). N = 3 to 4 mice per timepoint. Red bars: mean. D) Representative immunofluorescence images showing localization of neutrophils (Ly6G^+^, top row) and macrophages (CD68^+^, bottom row) around a stage 3 SAC. Right column: zoom of white boxed area. Statistics: C) Kruskal-Wallis one-way ANOVA with Dunn’s post-test. \**p*<0.05, n.s.: not significant.

Many stage 4 abscesses also had host cells interacting with small clusters of *S. aureus* near the SAC, possibly indicating phagocytic uptake of bacteria that were previously part of the SAC (**Supp Fig 1A**). Although stages 3 and 4 had significantly larger areas than stage 2, we did not observe differences within a given stage across days (**Fig 2C**). Multiple stages could typically be observed within a single kidney across timepoints, but the proportion of stages shifted over time (day 3 vs. day 5).

Immunofluorescence microscopy was used to visualize the localization of neutrophils (Ly6G^+^) and macrophages (CD68^+^) relative to *S. aureus*. We observed neutrophils near *S. aureus* at each stage, and stage 3 and 4 SACs were surrounded by large numbers of neutrophils (**Fig 2D, top row; Supp Fig 1B, right**). Although macrophages were also found near stage 1 and 2 bacteria, they were present at much lower numbers than neutrophils (**Supp Fig 1B, left**). There was a characteristic macrophage ring observed around stage 3 and 4 abscesses outside the neutrophil layer, away from the SAC (**Fig 2D, bottom row**) (26). This indicates that although macrophages are present within abscesses, *S. aureus* is predominantly interacting with neutrophils.

### Characterizing Agr and Sae fluorescent reporter activity in culture

Before completing infections with fluorescent reporter strains, we confirmed the *agr* and *sae* reporter strains and GFP^-^ control strain exhibited similar growth kinetics in liquid media (**Supp Fig 2A**). The *agr* reporter strain (*P_agrB_::gfp*) demonstrated a growth-dependent increase in reporter expression (GFP/mCherry ratio) as expected, since Agr has heightened activity at high cell density (**Supp Fig 2B**) (13). The *sae* reporter strain (*P_saeP_::gfp*) showed low expression in exponential phase as observed in previous studies (**Supp Fig 2B**) (27), and an increase in the percentage and mean fluorescence intensity of *sae-*expressing cells in late stationary phase (between 8h-24h culture) (**Supp Fig 2B-E**). Colonies of the GFP^-^ control strain had very low GFP fluorescence, while *agr* and *sae* reporter strains had high and intermediate levels of GFP fluorescence, respectively (**Supp Fig 2F**).

### *agr* reporter expression is not above baseline during kidney abscess development

To understand how the activity of the Agr and Sae systems changes during abscess development, we infected mice with the GFP^-^ control, *agr* (*P_agrB_::gfp*), or *sae* (*P_saeP_::gfp*) reporter strains. Kidneys were harvested at days 3, 4 and 5 p.i., corresponding with periods of peak bacterial growth within the kidneys. We did not detect significant differences in bacterial load, except a slight decrease in the CFUs of *sae* reporter strain at day 5 p.i. (**Supp Fig 3A**).

To quantify reporter expression, the constitutive mCherry signal was used to identify bacterial cells, and the total GFP (reporter) and mCherry signals were expressed as a GFP/mCherry ratio. The number of events per stage analyzed for each mouse is summarized in **Supplemental Table 1**. The GFP/mCherry ratio of *S. aureus* GFP^-^ control cells revealed variability across stages, within a stage across days, and even within a stage on a specific day (**Supp Fig 3B**). This could arise from leaky expression of GFP, differences in tissue autofluorescence, or differences in mCherry expression, the latter of which could be driven by changes in bacterial metabolic state. To take this inherent variability into account, the GFP/mCherry ratio of each event (belonging to a specific stage and day) was normalized to the average GFP^-^ control value at the same stage and day (black dashed baseline at Y=1) (**Supp Fig 3C**). A comparison of *agr* and *sae* reporter strain values before and after normalization to the GFP^-^ control strain is shown in **Supp Fig 3D-G**.

We first analyzed *agr* reporter expression within intracellular *S. aureus* (Stage 1), and found the average GFP/mCherry ratios were generally below the GFP^-^ control baseline indicating that the Agr system is off (**Fig 3A, 3B, Supp Fig 3E**). In stages 2-4, we observed a trend of increased relative *agr* expression at day 5 compared to day 3 (**Fig 3C**). A comparison with the GFP^-^ control strain indicated the *agr* GFP/mCherry ratio was statistically above baseline only in intact SACs (stage 3) at day 5 p.i. (**Supp Fig 3E**). Since Agr is a quorum sensing system and its activation is driven by *S. aureus* population density, we hypothesized the increased values would correlate with increased SAC size. However, we observed no significant differences in the size of a specific stage across days (**Fig 3D**). Within the stage 3, day 5 p.i. SACs, a majority had GFP/mCherry ratios similar to the GFP^-^ control (Low), while a few displayed values above the GFP^-^ control (High) (**Fig 3E, F**). We observed decreased mCherry signal (**Fig 3E**) in some SACs, which could result in lower ratiometric values, independent of changes in GFP signal. To understand which signal drove changes in GFP/mCherry ratios, we analyzed the signals separately, and found no significant changes in either GFP or mCherry signal when compared across groups (**Fig 3G**). These data indicate that decreased mCherry signal can raise the ratiometric value of individual SACs, and that *agr* expression was not above baseline during kidney abscess development.

**Figure 3:**
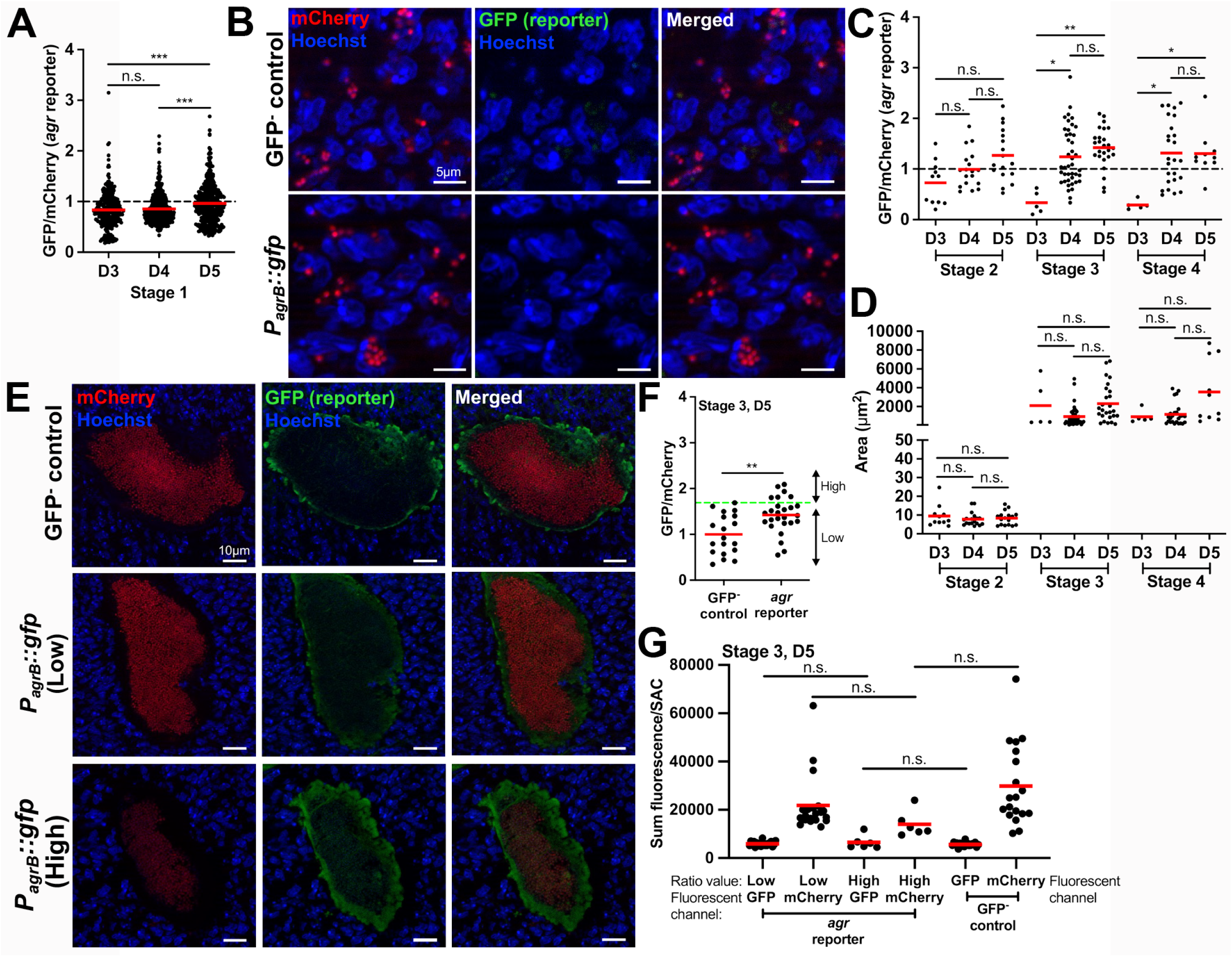
a*g*r reporter expression is not above baseline during kidney abscess development. C57BL/6 mice were inoculated with GFP^-^ control or *agr* reporter (*P_agrB_::gfp*) strains, both expressing mCherry constitutively. Mice were sacrificed at days 3, 4, or 5, and kidneys were harvested and processed for fluorescence microscopy. The GFP/mCherry ratio of each *agr* reporter event was normalized to the average value of the respective stage and day matched GFP^-^ control events (represented by the black dotted baseline at Y=1 in panels A and C). A) *agr* reporter expression in stage 1 *S. aureus*. Dots: intracellular *S. aureus* events. B) Representative images (D4) for data shown in panel A. C) *agr* reporter expression in stages 2 to 4. D) Size distribution of *agr* reporter extracellular clusters (stage 2) or SACs (stages 3 and 4) depicted in panel C. E) Representative images for data shown in panel F. F) Comparison of fluorescent signals in GFP^-^ control and *agr* reporter stage 3 SACs at day 5 p.i. Green dashed horizontal line: highest GFP/mCherry value for the GFP^-^ control, used to differentiate reporter High and Low SACs for the *agr* reporter. G) Sum fluorescence/SAC of individual fluorescent channels (GFP, mCherry). *agr* reporter datasets are split based on the Low or High ratio designation in F. All SACs from F are analyzed in G. Dots: individual events in panels C, D, F, and G. N = 3 to 4 mice per timepoint. Red bars: mean. Statistics: A), C), D), G) Kruskal-Wallis one-way ANOVA with Dunn’s post-test; F) Mann-Whitney. \*\*\**p*<0.001, \*\**p*<0.01, \**p*<0.05, n.s.: not significant.

### *sae* reporter expression is highly heterogenous in intracellular *S. aureus* but is more uniformly high within intact abscesses

We next analyzed *sae* reporter expression during abscess development. Unlike the *agr* reporter, the average values of the *sae* reporter stage 1 events were higher than baseline on days 3 and 5 p.i. suggesting that the Sae system is generally active during intracellular residence of *S. aureus* within mouse kidneys (**Fig 4A**). Interestingly, we observed a subset of stage 1 *S. aureus* exhibited *sae* expression levels (GFP/mCherry) similar to the GFP^-^ control (Low), while other bacteria had higher expression levels (High, 18.7% of the population) (**Fig 4B**). Analysis of individual fluorescent channels indicated that the GFP reporter signal was significantly above baseline for ‘High’ cells without a significant drop in mCherry (compared to the GFP^-^ control), confirming these cells were *sae*-expressing (**Fig 4C**). We also observed that neutrophils (Ly6G^+^) harbor both subsets of *sae*-Low and *sae-*High *S. aureus* (**Fig 4D**). At stage 2, *sae* expression was only above baseline on day 3. Within each day, there was an increase in *sae* expression at stages 3 and 4, suggesting the Sae system is highly active within intact and dispersed SACs (**Fig 5A, 5B**).

**Figure 4:**
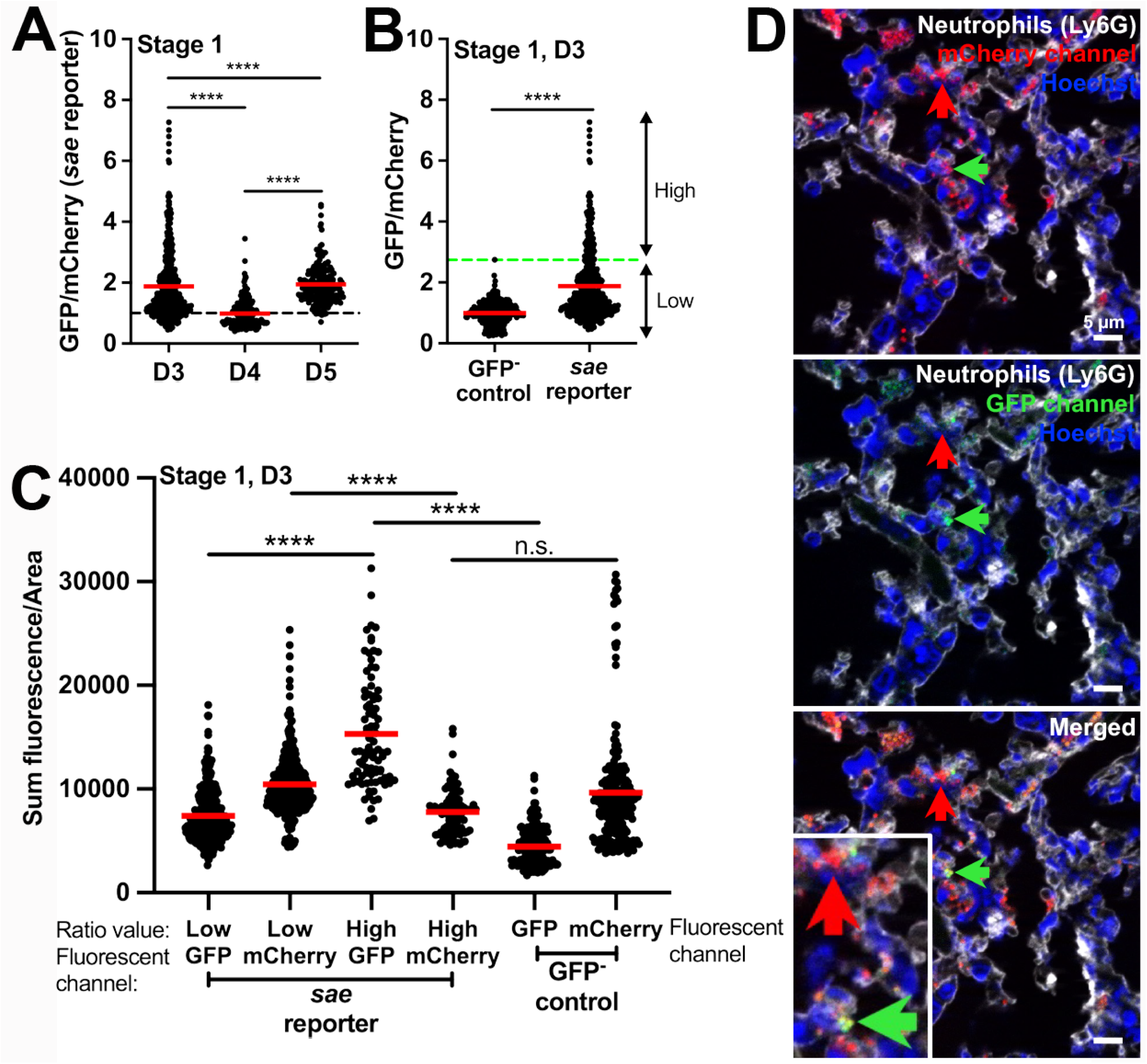
s*a*e reporter expression is highly heterogenous in intracellular *S. aureus*. C57BL/6 mice were inoculated with GFP^-^ control or *sae* reporter (*P_saeP_::gfp*) strains, both expressing mCherry constitutively. Mice were sacrificed at days 3, 4, or 5, and kidneys were harvested and processed for fluorescence microscopy. The GFP/mCherry ratio of each *sae* reporter event was normalized to the average value of GFP/mCherry ratios of the respective stage and day matched GFP^-^ control events (represented by the black dotted baseline at Y=1 in panels A and B). A) *sae* reporter expression in stage 1 *S. aureus*. Dots represent intracellular events. B) Comparison of fluorescent signals in GFP^-^ control and *sae* reporter stage 1 events at day 3 p.i. Green dashed horizontal line: highest GFP/mCherry value for the GFP^-^ control, used to differentiate reporter Low and High events for the *sae* reporter. C) Sum fluorescence/Area of individual fluorescent channels (GFP, mCherry). *sae* reporter datasets are split based on the Low or High ratio designation in B. All SACs from B are analyzed in C. D) Representative images for data shown in panel B. Red arrow represents *sae* non-expressing (Low) *S. aureus*, green arrow represents *sae*-expressing (High). Inset on merged image is a zoom-in of highlighted Low and High cells. N = 3 to 5 mice per timepoint. Red bars: mean. Statistics: A) and C) Kruskal-Wallis one-way ANOVA with Dunn’s post-test; B) Mann-Whitney \*\*\*\**p*<0.0001, n.s.: not significant.

**Figure 5:**
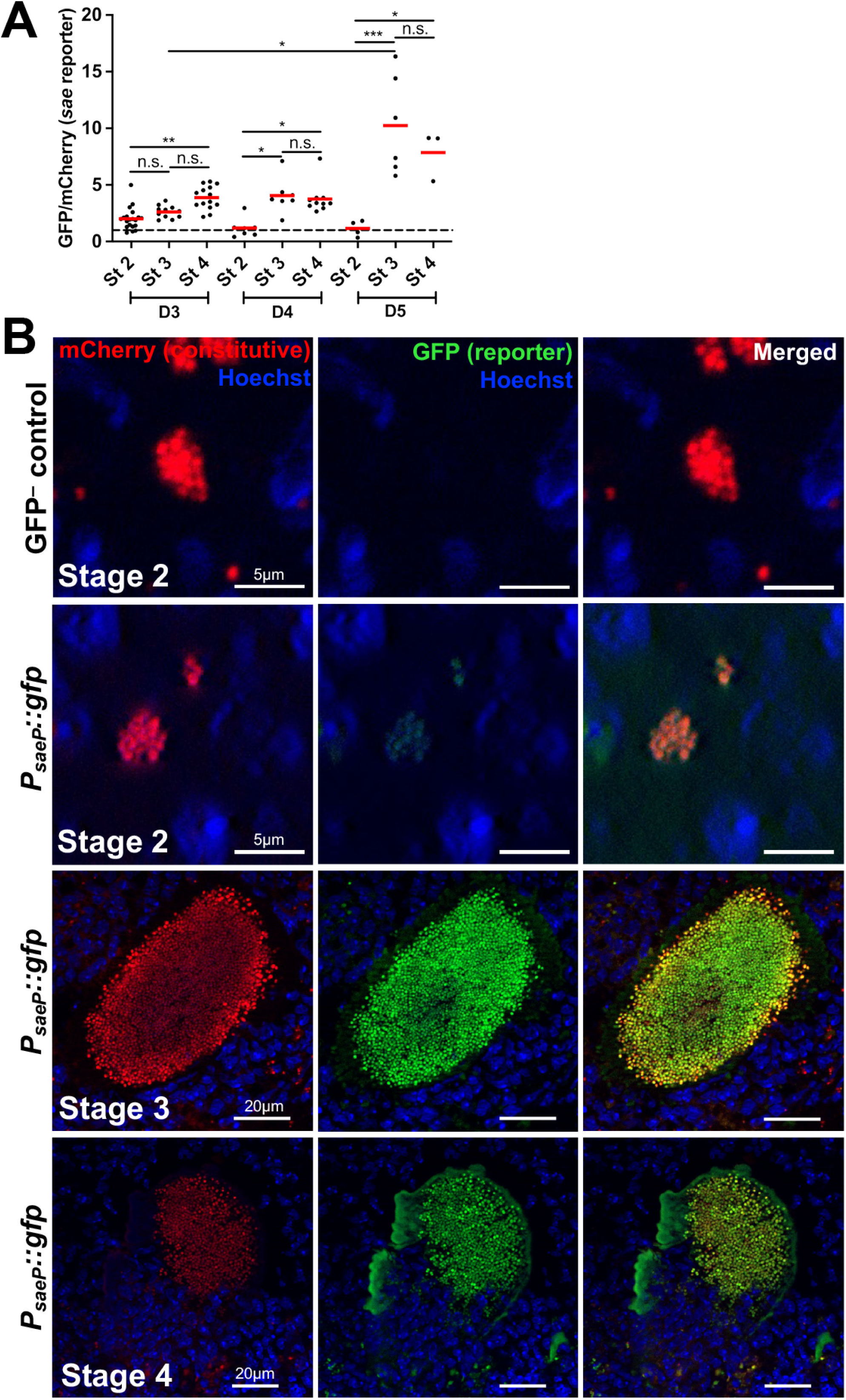
s*a*e reporter expression is higher in fully formed SACs as compared to small extracellular clusters. C57BL/6 mice were inoculated with GFP^-^ control or *sae* reporter strains, both expressing mCherry constitutively. Mice were sacrificed at days 3, 4, or 5, and kidneys were harvested and processed for fluorescence microscopy. The GFP/mCherry ratio of each *sae* reporter event was normalized to the average value of GFP^-^ control events (represented by the black dotted baseline at Y=1 in panel A). A) *sae* reporter expression in abscess stages 2 to 4. Dots represent individual events. N = 3 to 5 mice per timepoint. Red bars: mean. B) Representative images. Statistics: A) Kruskal-Wallis one-way ANOVA with Dunn’s post-test. \*\*\**p*<0.001, \*\**p*<0.01, \**p*<0.05, n.s.: not significant.

### The *sae* reporter exhibits heterogeneity in spatial expression patterns across SACs

We hypothesized that peripheral bacteria within SACs (stages 3 and 4) may experience a different microenvironment than bacteria at the SAC center due to immune cell interactions and higher concentrations of nutrients. Since these signals could modulate Sae activity, we sought to determine if there were spatial differences in *sae* expression within individual SACs. We observed 3 categories of SACs: 15/22 exhibited a Center^high^ Periphery^low^ phenotype (center to periphery >1.0), 5/22 had lower expression at the center (Center^low^ Periphery^high^, center to periphery <1.0), and 2/22 had similar expression at the center and periphery (center to periphery =1.0) (**Fig 6A, 6B**). In dispersed stage 4 SACs, we compared *sae* expression in peripheral *S. aureus* that were ‘in contact with host cells’ (along the point of rupture) with bacteria embedded deep within the SAC (‘away from host cells’) (**Fig 6C, 6D**). Here we also observed 3 phenotypes: 11/26 with higher expression away from host cells (>1.0), 8/26 with lower expression away from host cells (<1.0), and 7/26 with equivalent expression at both locations (=1.0). However, differences were modest (range: 0.52-1.28) (**Fig 6C**). The GFP and mCherry values at the center and periphery for each SAC are shown in **Supp Figs 4A-B**. Collectively, these data indicate that *sae* reporter expression is highly heterogeneous between individual abscesses.

**Figure 6:**
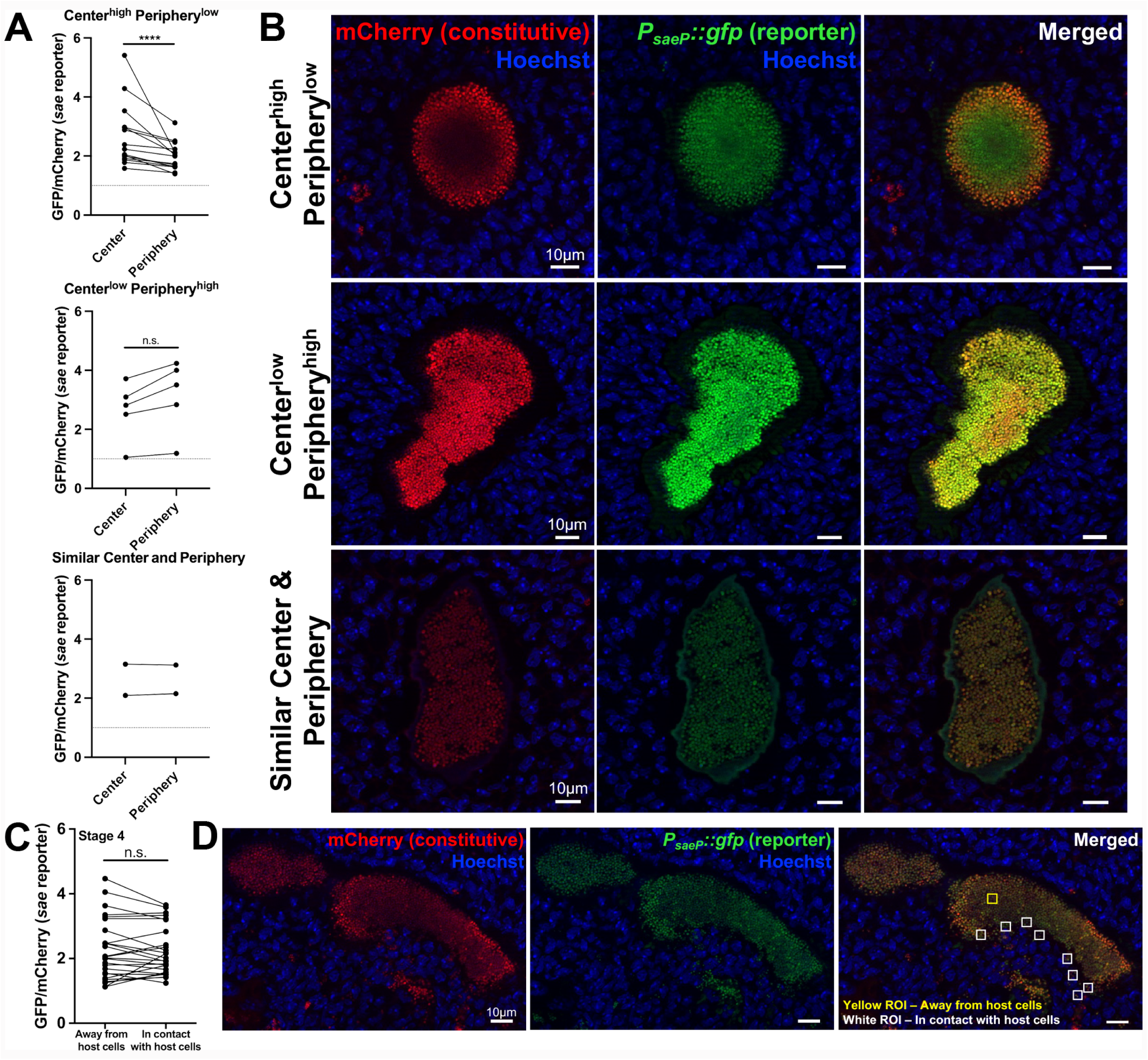
The *sae* reporter exhibits heterogeneity in spatial expression patterns across SACs. C57BL/6 mice were inoculated with the *sae* reporter strain. Mice were sacrificed at days 3, 4, or 5, kidneys were harvested and processed for fluorescence microscopy. Stage 3 (A, B): one region of interest (ROI, 4µm^2^) at the center and 8 ROIs along the periphery were selected. The GFP/mCherry ratio for each ROI was calculated using sum GFP and sum mCherry values. The GFP/mCherry ratio of the center and the average GFP/mCherry value of the 8 peripheral ROIs are shown. Stage 4 (panel C,D): one ROI in the center (away from host cells) and 8 ROIs along the rupture (in contact with host cells) were selected. The GFP/mCherry ratio at the center versus averaged periphery of A) stage 3, combined data from D3 to D5, split by spatial phenotype; and C) stage 4, combined data from D3 to D5. Lines connect values from the same SAC. Representative images are shown in B and D. N = 3 to 5 mice per timepoint. Statistics: A) and C) Wilcoxon matched-pairs test. \**p*<0.05, n.s.: not significant.

### Loss of *sae* but not *agr* impacts abscess development

Given our data that Agr is predominantly inactive and Sae is active during kidney abscess development, we hypothesized that loss of *sae*, but not *agr*, would hinder abscess development. To test this, we performed infections with WT, *agr::tet* (28) and *saeQRS::spec* (29) strains constitutively expressing mCherry. Day 1 timepoints were used to assess relative amounts of bacterial seeding in the kidney, and day 4 was used to represent the peak of infection. CFU quantification indicated that the WT bacterial load increased between day 1 and day 4 p.i., but the *saeQRS::spec* CFUs did not increase (**Fig 7A**). To investigate abscess development, we performed microscopic analysis at day 4 p.i., where bacteria in 2 distinct sections were imaged from each tissue and categorized into abscess stages. Intracellular *S. aureus* (stage 1) and small extracellular clusters (stage 2) were observed in mice infected with each strain (**Fig 7B, 7D**). However, major differences were detected in the ability of the *saeQRS::spec* strain to progress to later stage SACs. Intact (stage 3) and dispersed (stage 4) SACs were observed for the WT and *agr::tet* strains, but we did not detect any stage 3 SACs for the *saeQRS::spec* strain, and observed a single, small *saeQRS::spec* stage 4 SAC with aberrant morphology relative to the WT strain (**Fig 7C, 7E**). Our data suggest that an active Sae system is necessary for the formation of SACs and kidney abscess development, while Agr is dispensable.

**Figure 7:**
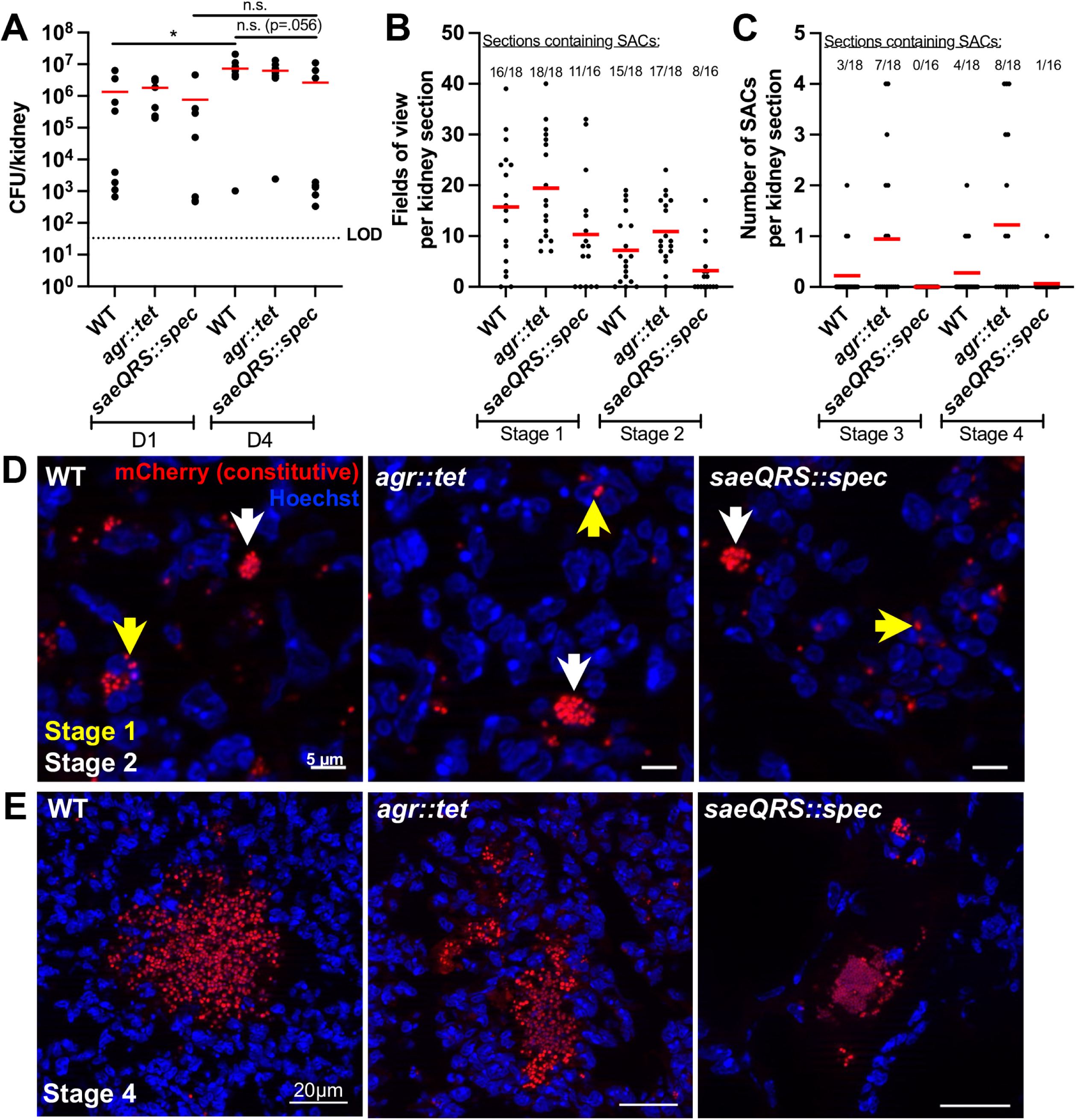
Loss of *sae* but not *agr* impacts abscess development. C57BL/6 mice were inoculated with WT, *agr::tet*, or *saeQRS::spec* strains expressing mCherry constitutively. Mice were sacrificed at days 1 or 4 and kidneys were harvested. Left kidneys were homogenized to quantify bacterial load (CFU/kidney) and right kidneys were fixed and processed for fluorescence microscopy. (A) Bacterial load, each dot represents one mouse. N = 7 to 9 mice per timepoint. Red bars: mean. Dotted line denotes the limit of detection (LOD). (B) Number of fields of view per section that contained stage 1 or stage 2 SACs (D4 p.i.). N = 8 to 9 mice (2 sections per mouse). (C) Number of stage 3 or stage 4 SACs per section (D4 p.i.). N = 8 to 9 mice (2 sections per mouse). (D) Representative images for data shown in panel B. Yellow and white arrows represent stage 1 and stage 2 events, respectively. (E) Representative images for data shown in panel C. Statistics: A) Kruskal-Wallis one-way ANOVA with Dunn’s post-test. \**p*<0.05, n.s.: not significant.

## Discussion

The Agr and Sae master regulatory systems control the expression of a large arsenal of *S. aureus* virulence factors, including those that contribute to kidney abscess development (9, 17). However, when they are active, and if activity is restricted to specific bacterial subpopulations, has remained largely unknown. To address this question, we first generated a modified categorization to define the stages of kidney abscess formation (**Fig 2A**). Our staging was very similar to Cheng et al. (24), but we focused on events in the kidney omitting the initial bloodstream survival stage (9, 24). We also utilized the USA300 LAC strain, which unlike strain Newman (15, 30), has the ability to regulate Sae. Somewhat surprisingly, we observed a great deal of similarity in abscess development compared to strain Newman (9), which may be linked to our observation that Sae is largely during abscess development. A notable exception was observed at early stages, when intracellular *S. aureus* within neutrophils exhibit a clear ON/OFF Sae phenotype (**Fig 4B, 4C**). Given *sae* expression is required for abscess progression (**Fig 7B, 7C**), this could indicate ON and OFF cells have different intracellular fates. Alternatively, this could indicate that some cross-complementation occurs, where Sae activity in a subset of cells may be sufficient to modulate the host environment and promote the growth and progression of the bacterial population as a whole.

*sae* expression increased within mature SACs, and we observed 3 distinct *sae* expression patterns (**Fig 6**). The set with higher *sae* expression at the periphery was consistent with the model that immune cell proximity induces heightened Sae activity. SACs with heightened *sae* expression at the center were instead consistent with Behera et al., where higher expression of Sae-regulated *nuc* occurred at the SAC center (31).

Nutrient depletion can relieve CodY repression of *sae* expression (16). Consistent with this, we observed an increase in *sae* reporter expression in the extended stationary phase of culture (**Supp Fig 1C-E**). Differential nutrient availability is known to promote heterogeneity in other bacterial abscess and granuloma models (32, 33), and it is possible that bacteria at the center of SACs experience nutrient limitation, triggering *sae* expression. It is also likely that individual abscesses have distinct nutritional microenvironments, given the *S. aureus* inter-abscess heterogeneity in metal availability and siderophore production (34, 35). We would predict that the heterogeneity we observed is a cumulative effect of these factors.

While we observed the importance of Sae for abscess development, we saw surprisingly little Agr activity and found that *agr* was not required for kidney abscess progression. A previous study by García-Betancur et al. showed that downregulation of Agr activity can be driven by Mg^2+^, and Mg^2+^-rich organs like kidneys and bones were found to harbor higher proportions of Agr-inactive *S. aureus* cells (36). Additionally, the macrophage apolipoprotein B receptor (*Apobr*) is known to inhibit the Agr system, and its presence has been detected at the SAC-host interface within kidney abscesses (37). However, using a 3D *in vitro* model of SACs, Guggenberger et al. showed that Agr is required for the dispersion of SACs at later stages (25). Given dispersal was observed with the *agr* mutant strain, additional experiments would be needed to determine the contribution of Agr to SAC dispersal in the kidney.

A key limitation of these findings is that they were restricted to endpoint analyses.

Tracking individual abscesses overtime could yield even more novel spatiotemporal information regarding virulence factor expression and abscess biology. Use of a luciferase reporter with IVIS technology can enable real-time monitoring of gene expression during infection, but due to limitations in resolution, spatial gene expression information cannot yet be obtained (5). Two recent studies have utilized 3D tissue

culture-based SAC models that could potentially enable spatiotemporal monitoring (25, 38). These systems also offer the ability to incorporate human components such as fibrinogen, immune cells, or tissue-specific cells to facilitate translation of findings to human infection. Data obtained from these diverse but complimentary model systems can help generate an improved picture of *S. aureus* pathogenesis and aid in the development of strategies for better management of staphylococcal infections.

## Materials and Methods

*Bacterial strains & growth conditions. S. aureus* USA300 LAC wildtype (39) and its derivatives were grown in tryptic soy broth (TSB), and *E. coli* DH5α and IM08B were grown in Luria broth (LB), all at 37°C with aeration. Antibiotics were used at the following concentrations: kanamycin (50µg/ml), tetracycline (4µg/ml), and spectinomycin (250µg/ml) for *S. aureus*; erthyromycin (500µg/ml) and chloramphenicol (25µg/ml) for *E. coli*.

*Construction of S. aureus fluorescent reporter strains.* To generate constitutive mCherry^+^ strains, the *P_sarA_*::*sod*RBS fragment from pOS1-*P_sarA_*-*sod*RBS-*sgfp* (29) was fused to a *S. aureus* codon-optimized *mCherry* (IDT) and cloned into pJC1111 for chromosomal integration at the SaPI1 *attC* attachment site (20). The plasmid was isolated from *E. coli*, electroporated into *S. aureus* strain RN9011 (20), recombinants were selected on 0.1 mM cadmium chloride, and the plasmid was transferred into strain LAC by phage transduction.

For GFP reporters, pIMAY, a temperature-sensitive plasmid with a tetracycline-inducible anti-*secY* counter selection marker, was used (40). For the GFP^-^ control strain, pINT-*gfp* was constructed by assembling in pIMAY: *sod*RBS-*sgfp*, *kanR* cassette, and flanking homology arms for genome integration at a neutral site between SAUSA300_RS05730 and RS05735 (21). To construct pINT-*P_agrB_*::*gfp* and pINT-*P_saeP_*::*gfp*, promoter regions upstream of *agrB* and *saeP* were cloned into pINT-*gfp*. Plasmids were assembled and isolated from *E. coli* IM08B, then transformed directly into *S. aureus* LAC strains (41).

For strains containing both fluorescent constructs, the GFP construct was added first. Transformants were selected on chloramphenicol at 28°C, chromosomal integration was facilitated by growth at 37°C, and recombinants were selected on 1 µg/ml anhydrotetracycline, as described in (40). Additional reporter construction details are included in the Supplemental Information file.

*In vitro reporter characterization.* Overnight cultures (16h) were diluted 1:100 into fresh TSB and incubated at 37°C with aeration. At each timepoint, samples were added to black-walled, clear bottom, 96-well plates for absorbance (OD_600nm_) and fluorescence measurements (GFP: 480ex/520em, mCherry: 560ex/610em) using a Synergy microplate reader (Biotek Instruments). Colonies on tryptic soy agar were imaged using a Biotek Cytation 5 cell imaging multimode reader (Agilent). For flow cytometry, 100 µl of the culture at the indicated OD was pelleted and fixed in 4% paraformaldehyde (PFA) in PBS overnight at 4°C. Samples were analyzed on a BD FACSymphony A3 flow cytometer. Data was analyzed using FlowJo v10.10.0 software (BD Biosciences).

*Murine infection model.* All animal experiments were approved by the Johns Hopkins University Institutional Animal Care and Use Committee (Protocol #s: MO20H330, MO23H310). *S. aureus* frozen stocks in 10% glycerol (prepared after 2h of growth from a 1:50 back-dilution of overnight cultures) were washed and diluted in sterile PBS to 10^6^ CFU in 100 µl for inocula. 6-to 8-week-old female C57BL/6 mice (Jackson Laboratories) were inoculated intravenously via the retro-orbital (under isoflurane anesthesia) or tail vein (with manual restraint) routes. At the indicated timepoints post-inoculation, mice were euthanized using a lethal dose of isoflurane followed by cervical dislocation, and kidneys were harvested. Left kidneys were homogenized and used for CFU enumeration and the right kidneys were processed for microscopy after fixation in 4% PFA overnight at 4° C (exception: Figure 1B, left/right CFU comparison).

*Fluorescence microscopy of mouse tissues.* After fixation, the right kidneys were frozen-embedded in O.C.T. compound (Tissue-Tek, VWR) and stored at-80°C. 10 µm sections were cut using a cryostat microtome (Microm HM 505e) and mounted on charged microscope slides (HistoBond, VWR). Sections were thawed in PBS at room temperature (RT) and stained with Hoechst (1:10,000 dilution in PBS) for 15 minutes.

To stain immune cells, thawed sections were permeabilized using ice-cold methanol for 2 min, blocked with 2% bovine serum albumin (BSA) in PBS at RT for 1h, then incubated with either APC-conjugated rat anti-mouse Ly6G antibody (Invitrogen: 17-9668-80) or rat anti-mouse CD68 primary antibody (BioRad: MCA1957T) diluted in 2% BSA overnight at 4°C. For CD68 detection, sections were washed in PBS and incubated with donkey anti-rat Alexa Fluor 647-conjugated secondary antibody (Invitrogen:

A48272TR) for 1h at RT and stained with Hoechst. Coverslips were mounted with ProLong Gold (Invitrogen). 2-3 sections per mouse were imaged using a Zeiss Axio Observer 7 inverted fluorescent microscope with a 63x oil objective and Apotome.2. Images were captured with an Axiocam 702 mono camera (Zeiss) and processed using ZEN3.10 software.

### Image Analysis

#### Abscess stage criteria

Stage 1: Intracellular *S. aureus* (single cells or clusters) in contact with host cell nuclei (detected with Hoechst).

Stage 2: Small extracellular clusters (distanced from host cells).

Stage 3: Large compact or loosely-packed SACs, surrounded by a continuous fibrin layer, with no direct host cell contact.

Stage 4: Dispersed SAC with a discontinuous fibrin layer and host cells contacting *S. aureus* at multiple points.

Stage 1 and 2 events were excluded if found in the same field as stage 3 or 4 SACs, to ensure these events represented early stages preceding SAC formation, and not ruptured SACs.

#### Temporal expression pattern

Volocity image analysis software was used to quantify bacterial area, mCherry, and GFP signals. Objects (individual bacterial cells/cluster/SACs) were defined by mCherry signal, and area, sum mCherry, and sum GFP fluorescence were measured. Sum GFP/sum mCherry ratios were calculated.

Objects <0.4 µm^2^ (artifacts) were excluded, but faint mCherry^+^ cells were included. For stage 1, at least 100 events or events from 2 fields per section (whichever was greater) were analyzed. For stages 2-4, all events per section were analyzed to avoid selection bias. If <5 events were present, an additional section (separated by ≥150µm) was analyzed to reach 5 events per mouse.

#### Spatial expression pattern

For stage 3 SACs, one region of interest (ROI: 4×4 µm) at the center and 8 ROIs at the periphery were selected. For stage 4 SACs, one interior ROI (away from host cells) and 8 ROIs along rupture sites (in direct contact with host cells) were selected. Within these ROIs, objects were detected as described above, and the GFP/mCherry ratio for each ROI was calculated using sum GFP and sum mCherry values. The GFP/mCherry ratio of the center and the average GFP/mCherry value of the 8 peripheral (or in direct contact) ROIs are shown.

#### Comparison of abscess stage distribution in WT, agr::tet and saeQRS::spec infections

2 sections approximately 300 µm apart were analyzed per mouse. All bacteria-containing fields were imaged as described above. Due to the high abundance of stages 1 and 2, the number of fields containing at least one stage 1 or 2 event were counted per section. For stages 3 and 4, the numbers of SACs per section were quantified.

#### Statistical analysis

GraphPad Prism 10 was used to graph all data and for statistical analyses. The statistical test performed for each experiment is described in the respective figure legend. A *P* value of < 0.05 was considered significant.

## Author Contributions

Conceptualization: VJT, KMD; Formal Analysis: AA, REB, KMD; Funding Acquisition and Supervision: KMD; Investigation: AA, REB, BL, VA, CA, AW, ALE, MHC, KMD; Methodology: BL, II, RP; Writing – Original Draft Preparation: AA, REB; Writing – Review & Editing: AA, REB, ALE, MHC, II, VJT, KMD.

## Acknowledgments

We thank the Davis, Archer, Klein, Pekosz, Thompson and Baumgarth labs for constructive feedback and suggestions. Special thanks to Parsa Farhang (Davis lab) for manuscript editing support. We also thank the JHSPH Flow cytometry Core and MMI Microscopy facilities. This work was supported by NIH/NIAID AI154116 and JHSPH Faculty Innovation Fund grants to KMD and NIH/NIAID AI105129 and AI099394 to VJT. The funders had no role in study design, data collection and interpretation, or the decision submit the work for publication.

## Supporting Information

**Supplemental Figure 1: Immune cell localization in mouse kidney abscesses.** C57BL/6 mice were inoculated with the *S. aureus* GFP^-^ control strain. Mice were sacrificed at day 4 and kidneys were harvested and processed for fluorescence microscopy. (A) Representative image showing localization of neutrophils (Ly6G^+^) around a dispersed stage 4 SAC. White arrows: *S. aureus* interacting with neutrophils in the vicinity of a dispersed SAC. (B) Representative image showing localization of macrophages (CD68^+^, left panel) and neutrophils (Ly6G^+^, right panel) around stage 1 and stage 2 events.

Supplemental Figure 2: Characterizing *S. aureus* fluorescent reporter strains during growth *in vitro* in tryptic soy broth or agar. Overnight cultures of *S. aureus* were diluted 1:100 in fresh TSB and incubated at 37°C with shaking. At the indicated timepoints, (A) absorbance (OD_600nm_), (B) GFP, and mCherry fluorescence were measured using a microplate reader. Mean ± SD of three biological replicates are shown. Black dotted line (baseline): average GFP/mCherry value of the GFP^-^ control at the indicated timepoints. (C) Representative flow cytometry plot showing reporter expression in GFP^-^ control and *sae* reporter strains at the indicated growth phases and timepoints. (D) Percentage of GFP^+^ *sae* reporter cells at the indicated timepoints, measured by flow cytometry. (E) Mean fluorescence intensity (MFI) of GFP in the *sae* reporter strain. Mean values represent 50,000 cells per replicate, N = 3 biological replicates. Lines connect values from the same replicate. (F) Colonies of *S. aureus* on tryptic soy agar (kanamycin 50 µg/ml) imaged after overnight growth at 37°C. Statistics: B) Two-way ANOVA with Tukey’s test, comparison to GFP^-^ control. \*\*\*\**p*<0.0001, \*\**p*<0.01, \**p*<0.05, n.s.: not significant.

**Supplemental Figure 3: Reporter expression in mouse kidney abscesses.** C57BL/6 mice were inoculated with GFP^-^ control, *agr*, or *sae* reporter strains. Mice were sacrificed at days 3, 4, or 5 (D3, D4, D5) and kidneys were harvested. Left kidneys were homogenized to quantify bacterial load (CFU/kidney) and right kidneys were fixed and processed for fluorescence microscopy. (A) CFU/kidney of mice infected with *S. aureus* reporter strains at the indicated timepoint. Each dot represents one mouse. N = 3 to 6 mice. (B) GFP/mCherry ratio of GFP^-^ control abscesses (stages 1 to 4) at the indicated timepoints. (C) GFP/mCherry ratio of individual GFP^-^ control events after normalization to the average value of day-and stage-matched GFP^-^ control events (represented by the black dashed baseline at Y=1 in panel C, E and G). (D - G) Comparison in reporter expression of *agr* or *sae* reporter events (stages 1 to 4) to day-and stage-matched GFP^-^ control events; GFP/mCherry ratios without normalization (panels D and F) and after normalization to the GFP^-^ control (panels E and G) are shown. Dots represent intracellular *S. aureus* (single/cluster, stage 1), individual extracellular clusters (stage 2) or SACs (stages 3 and 4) in panels B to G. N = 3 to 5 mice per timepoint. Red bars represent mean. Statistics: A), B), C), E) and G) Kruskal-Wallis one-way ANOVA with Dunn’s test. \*\*\*\**p*<0.0001, \*\*\**p*<0.001, \*\**p*<0.01, \**p*<0.05, n.s.: not significant.

**Supplemental Figure 4: GFP (reporter) and mCherry (constitutive) signals within individual SACs.** C57BL/6 mice were inoculated with the *sae* reporter strain. Mice were sacrificed at days 3, 4, or 5, and kidneys were harvested and processed for fluorescence microscopy. Stage 3 (panel A): one region of interest (ROI, 4µm^2^) at the center and 8 ROIs along the periphery were selected. The 8 peripheral ROI values were averaged. Stage 4 (panel B): one ROI in the center (away from host cells) and 8 ROIs along the rupture (in contact with host cells) were selected. The ‘in contact’ ROI values were averaged. Shown are sum GFP and sum mCherry values at the center and periphery of (A) stage 3, combined data from D3 to D5 (GFP/mCherry shown in Fig 6C); and (B) stage 4, combined data from D3 to D5 (GFP/mCherry shown in Fig 6E). N = 3 to 5 mice per timepoint. Statistics: Wilcoxon matched-pairs test. \*\*\*\**p*<0.0001.

**Supplemental Table 1: Sample size information for microscopy analyses.** C57BL/6 mice were inoculated with *S. aureus* GFP^-^ control, *agr*, or *sae* reporter strains. At the indicated timepoints, mice were sacrificed, and kidneys were harvested. Left kidneys were used for CFU enumeration, and right kidneys were processed for microscopic examination. N represents the number of mice. 2-3 sections per mice were imaged and analyzed. The number of events analyzed for each mouse are shown.

**Supplemental Data File.** All raw data from the manuscript is provided in this file. **Supplemental Information File:** File includes a pINT-*gfp* plasmid map, *agrB* and *saeP* promoter sequences included in fluorescent reporter constructs, and additional methodological details for *S. aureus* fluorescent reporter strain construction.

